# Missing what’s right under your nose: failed appetitive and aversive audio-olfactory conditioning in humans

**DOI:** 10.1101/2024.12.17.628856

**Authors:** N.S. Menger, B. Kotchoubey, K. Ohla, Y.G. Pavlov

**Affiliations:** Institute of Medical Psychology and Behavioural Neurobiology, University of Tübingen; Perception & Cognitive Neuroscience, Science & Research, dsm-firmenich, Satigny

**Author notes:** **Corresponding author** Nick S. Menger, Institute of Medical Psychology and Behavioural Neurobiology, University of Tübingen Silcherstraße 5, 72074 Tübingen.

**Keywords:** Appetitive conditioning, aversive conditioning, olfaction, audio, EEG, autonomic physiology

## Abstract

The comparison of physiological mechanisms underlying appetitive and aversive conditioning is often challenging due to the involvement of stimuli from different modalities with potentially disparate effective mechanisms (e.g., pain stimuli versus monetary rewards). The olfactory system offers a unique opportunity to examine both types of conditioning in humans, as isointense odors can serve as comparably pleasant and unpleasant stimuli. To study physiological and behavioral responses during appetitive and aversive learning, we employed odors as unconditioned stimuli (US) in a within-subjects design, measuring various conditioned physiological responses including skin conductance, heart rate, pulse wave amplitude, respiration, fear-potentiated startle, postauricular reflex, facial electromyography as well as event-related potentials, and auditory steady-state responses (ASSR) derived from electroencephalography. We conducted four experiments with a total of 95 participants, presenting three neutral sounds paired with either a pleasant odor, unpleasant odor, or odorless air. The first experiment involved uninstructed participants and frequency-modulated conditioned stimuli (CS) for ASSR analysis. In the second experiment, we omitted the frequency modulation and startle probe. The third experiment included pre-experiment instruction on CS-US contingencies, while the fourth employed a delayed conditioning paradigm in contrast to the other three experiments. Our results revealed differences between CS+ and CS-only in the fear-potentiated startle response in Experiment 3. No other effects were found. The minimal or absent learning effects observed across multiple peripheral and neural physiological measures may be attributed to the extra-thalamic nature of olfactory pathways and the subsequent difficulty in forming associations with auditory stimuli.

**Impact statement:** In a series of 4 experiments, we explored the neurophysiological differences between appetitive and aversive conditioning. Yet, none of the experiments showed effective conditioning. We hypothesize that the lack of learning effects is attributed to the inherent difficulty in forming associations between auditory and olfactory inputs.

## Introduction

In human research, comparing appetitive and aversive conditioning presents challenges due to the nature of the sensory modalities typically involved. Aversive conditioning studies often use stimuli such as loud sounds or electric shocks (Lonsdorf et al., 2017), which lack direct pleasant counterparts for appetitive conditioning. While monetary rewards are frequently used in appetitive conditioning studies, they are secondary reinforcers (Delgado et al., 2006). Visual stimuli also pose difficulties, as unpleasant and pleasant images can evoke emotions that differ in intensity and habituation rates (Bernet et al., 2008). This challenge can be addressed with odors which can be of opposing valence while being controlled in other stimulus features like intensity.

Compared to other modalities, few studies have examined olfactory conditioning. However, a large variety of measures has been used to operationalize the conditioned responses (CRs). Increased galvanic skin responses (GSR) or skin conductance responses (SCR) were found to aversive-, but not appetitive CS (Exner et al., 2021; Hermann et al., 2000; Stussi et al., 2018). Heart rate derived from the electrocardiogram (ECG) was only significantly higher in the aversive CS+ than CS-, while no differences were found between the appetitive CS+ and CS-. (Exner et al., 2021). Although measures such as the SCR and heart rate are the gold standard in conditioning studies, they seem to primarily capture changes in arousal, and it remains unclear whether they can differ between appetitive and aversive learning.

In contrast, the fear-potentiated startle (FPS) and postauricular reflex (PAR), where muscle activity below the eye and behind the ear is measured, respectively, are assumed to capture valence-specific differences. The FPS is found to be enhanced by stimuli associated with an aversive odor (Miltner et al., 1994), while the PAR response is higher for stimuli associated with a pleasant odor compared to an odorless condition (Stussi et al., 2018). These measures, therefore, may provide good differentiation in valence and may be valuable to compare appetitive and aversive conditioning independent of arousal.

One could argue that emotions have specific qualities that go beyond just positive or negative valence. For example, the disgust evoked by an unpleasant odor not only contrasts with the happiness caused by an appetitive event but also differs from other negative emotions such as fear of pain. Facial electromyography (EMG) recordings can potentially indicate these more specific emotions, as muscle activity of the levator labii was shown to differentiate between odors of different valence (Armstrong et al., 2007; Kato & Yagi, 1994). Olfactory conditioned responses in the corrugator or zygomaticus muscles may not be produced in a combined appetitive and aversive conditioning experiment (Hermann et al., 2000), however can indicate a CS-effect in an aversive conditioning only experiment (Flor et al., 2002; Hermann et al., 2002).

Neural measures of CR can provide valuable insights into the psychological processes accompanying appetitive and aversive CS. With its high temporal resolution, electroencephalography (EEG) is especially suitable for investigating their development over time. However, existing evidence is insufficient to provide a clear picture. Hermann et al. (2000) found a larger N1 in appetitive CS+ compared to CS− trials, but this was not the case for aversive CS+ trials. In contrast, Flor et al. (2002) found an effect of aversive CS+ on the N1. Furthermore, the contingent negative variation was modulated by the CS type in Flor et al. (2002), but not in Hermann et al. (2000). Hermann et al. (2000) further observed an interaction between electrode placement and CS type in the late positive complex during the aversive condition, which was not present in the appetitive condition. These findings suggest differences between aversive and appetitive CS, but further verification is required.

The primary aim of the current study was to investigate the differences in physiological responses between olfactory appetitive and aversive conditioning, with the goal of uncovering potentially distinct underlying mechanisms of these two types of learning. To achieve this, we employed an audio-olfactory conditioning paradigm in four successive, related experiments.

## Methods

### Experimental design

During the acquisition phase in Experiment 1 (Figure 1), participants were presented with a fixation cross for 2 s, then a 41.5 Hz-modulated sound of a vowel (CS) was played for 5s. After a jittered interval of 1-1.348 s, a startle probe occurred randomly in 20% of trials. Then, 8.5 s after the CS onset, the olfactory presentation period of 6 s started, during which the odor (US) was presented for 500 ms synchronized with an inhalation phase. This would either be a pleasant odor (US+appetitive), aversive odor (US+aversive) or odorless air (US-). If no inhalation was detected during this period, the odor was presented at the end of this period. 3 s after the odor presentation, a visual analogue scale (VAS) prompted the participants to rate the odor’s pleasantness and intensity. The inter-trial interval (ITI) following the VAS lasted so long until the total trial time amounted to 25 s. In Experiment 4, the US was not presented in sync with the breathing cycle, but was always presented 4.5 after the CS onset; that is, the CS and US overlapped for 500 ms.

**Figure 1.**
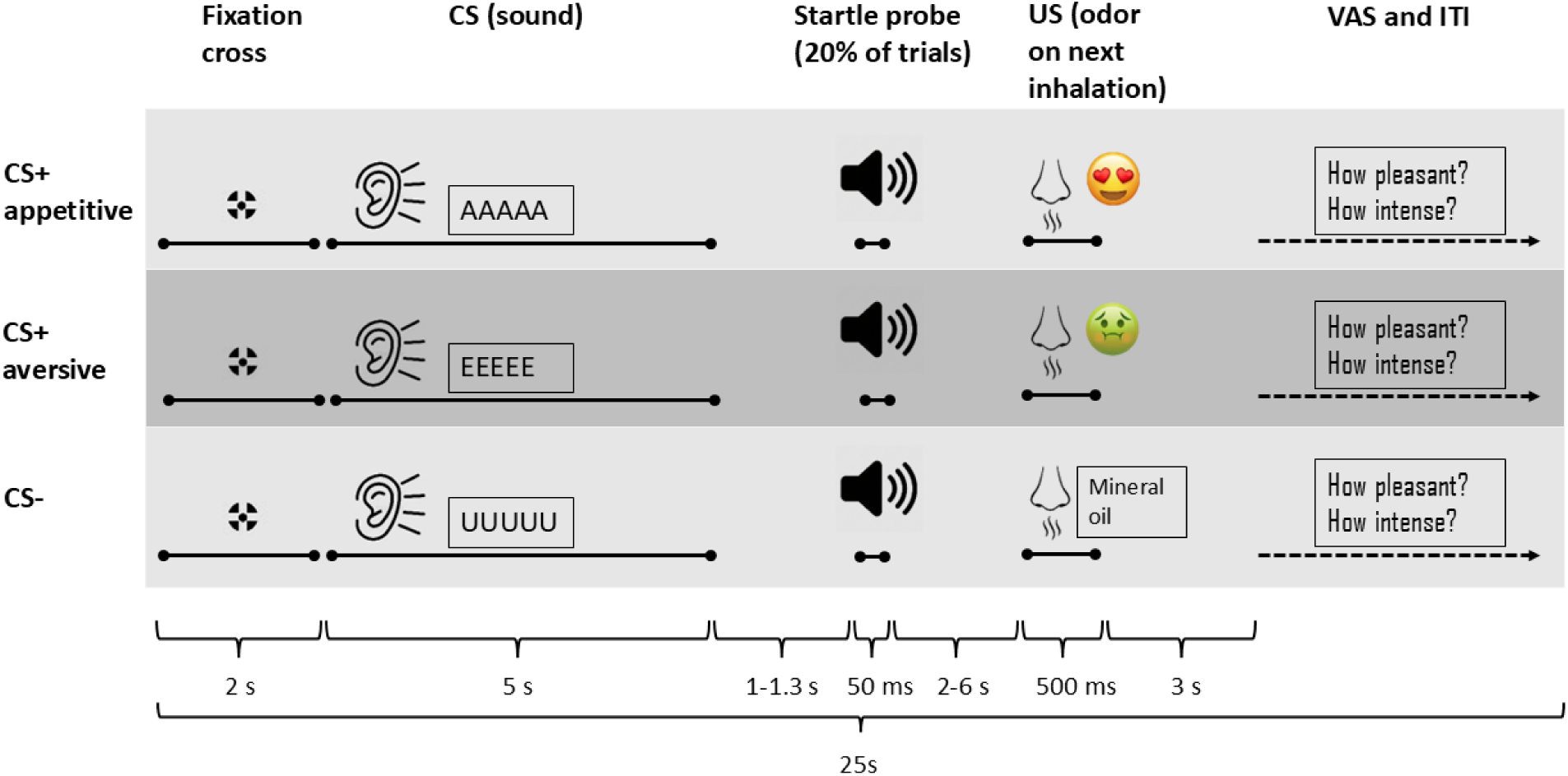
The experimental paradigm used in Experiment 1. At the start of the trial, the fixation cross would appear and remain on the screen until the VAS appeared at the end of the trial. Two seconds after the start of the trial, the CS was presented for five seconds, and after a jittered interval, the startle probe would occur in 20 % of the trials. Two seconds later, the olfactometer started tracking the participant’s respiration, and upon the next detected inhalation presented the olfactory US. Three seconds later, participants were prompted to rate the odor’s pleasantness and intensity, and would then wait during the ITI until the total trial time amounted to 25 s. Experiment 2 had a similar design as Experiment 1, but here the startle probe and sounds’ frequency modulations were omitted, and a 1 s interval after CS offset was used instead of the startle component. Experiment 3 had the exact same design as Experiment 1, with the exception that participants were instructed on the CS-US contingencies before the experiment. In Experiment 4, the startle probe and sound modulation were omitted, and the US was delayed to the CS, i.e. presented 4.5 s after the CS onset. The US was presented for 500 ms in all experiments.

We conducted four experiments, each differing in four parameters (Table 1). Experiment 2 was conducted to rule out the possibility that the CS frequency modulation and startle probes (which can be considered aversive and may decrease contingency awareness or conditioning effects; Sjouwerman et al., 2016) diminished conditioning that we observed in Experiment 1. These adjustments did not result in successful conditioning and only changed the awareness rate slightly. In Experiment 3, to further increase contingency awareness, participants were instructed on the CS-US contingencies, i.e. they were told which CS was followed by an appetitive US, aversive US, and US-. In addition, before the experiment, two training trials per condition were conducted to make sure that everyone would be fully aware of the contingencies. In Experiment 4, we used a delay conditioning design with overlapping CS and US, to test whether the long and variable interstimulus interval (ISI) employed in Experiments 1-3 led to unsuccessful conditioning.

**Table 1.** Differences in design for each experiment and corresponding awareness rates.

| Experiment | Startle probe | CS frequency modulation | Contingency instruction | Trace/delay conditioning | Awareness rate |
| --- | --- | --- | --- | --- | --- |
| 1 ($n=19$) | Yes | Yes | No | Trace | 31.58 % |
| 2 ($n=20$) | No | No | No | Trace | 50 % |
| 3 ($n=32$) | Yes | Yes | Yes | Trace | 100 % |
| 4 ($n=22$) | No | No | Yes | Delay | 100 % |

### Participants

Participants were recruited via email from the student and employee population of Eberhard Karls University of Tübingen for four experiments. The sample sizes and gender distributions were as follows: Experiment 1 (*n* = 21, 66.67 % female), Experiment 2 (*n* = 21, 54.54 % female), Experiment 3 (*n* = 36, 75 % female), and Experiment 4 (*n* = 24, 71.59 % female). The mean ages (and ranges) of participants were: Experiment 1 – 30.86 years (19 – 55), Experiment 2 – 30.18 years (19 – 54), Experiment 3 – 26.36 years (18 – 59), and Experiment 4 – 27.65 years (20 – 55). All participants were right-handed.

Exclusion criteria included psychological or neurological disorders, pregnancy, anosmia, respiratory disorders, and hearing impairments. Additional exclusions were necessary due to technical issues or inability to follow instructions: three participants from Experiments 1 and 2 (sound card problems), four from Experiment 3 (discomfort or technical issues with the EEG amplifier), and two from Experiment 4 (failure to report instructed contingencies). Furthermore, due to poor signal quality, a number of participants were excluded from the analyses of the different physiological measures. Final sample sizes used for each measure are reported in Tables 2 – 5.

Participants received either monetary compensation or course credit for their involvement. All participants provided informed consent prior to their involvement. The study adhered to the revised Declaration of Helsinki and received approval from the ethics board of the Medical Faculty at Eberhard Karls University of Tübingen.

The participants had a normal self-assessed sense of smell. On the day of the experiment, participants self-rated their smell ability on a scale of 1 – 100, where 0 corresponded to “no smell” and 100 to “excellent”. The mean ratings (and standard deviations) were: Experiment 1 – 76.7 (18.6), Experiment 2 – 72.4 (18.0), Experiment 3 – 71.2 (19.1), and Experiment 4 – 77.7 (19.9).

### Apparatus

Odorants were presented to participants’ nasal cavities using an olfactometer (Sniffo, CyNexo S.r.l.) via tubes connected to nose pieces inserted binasally approximately 0.5 cm deep. The odor channels had an airflow of 3 LPM for both nostrils combined, and were embedded in a constant airflow of 1 LPM. A fiber optic cable in one of the nose pieces measured the air temperature in the nasal cavity, determining the inhalation and exhalation phases of the respiration cycle.

Sounds were presented via an external soundcard (Focusrite Saffire Pro 14), using an amplifier (Atom Amp+, JDS Labs) and insert earphones (E-A-RTONE 3A, 3M). Participants used a gamepad (Logitech Gamepad F310) to rate stimuli on the VAS, using the D-pad to move the scale-slider and a button on the controller’s right side to confirm their choice.

The experiment took place in a soundproof, electrically shielded cabin containing a built-in monitor (LG Flatron L227WTP-PF) approximately 1 meter away from the participants’ eyes. The cabin’s lighting was set to 5 lux. Participants were seated in a comfortable chair for the duration of the experiment.

### Stimuli

Odorants were prepared as solutions in mineral oil, each with a total volume of 20 ml in 100 ml jars. The solutions were vortexed for 2.5 minutes using a Vortex-Genie 2 (Scientific Industries Inc.). Solute-solvent ratios were based on previous olfactory experiments (Amsellem et al., 2018; Höchenberger et al., 2015; Keller et al., 2012) and fine-tuned through a pilot study involving 14 participants who rated odor intensity and pleasantness. The following odorants were selected: Isoamyl acetate (1 % v/v; CAS 123-92-2), Orange oil (15 % v/v; CAS 8008-57-9), Lemon oil (5 % v/v; CAS 8008-56-8), Butyric acid (2 % v/v; CAS 107-92-6), Heptyl acetate (1 % v/v; CAS 112-06-1), Nonanal (5 % v/v; CAS 124-19-6), Linalool (1 % v/v; CAS 78-70-6), Mineral oil (100 % v/v; CAS 8042-47-5).

Sound stimuli consisted of 5-second utterances of the vowels A, E, and U, recorded in German by a female voice actor. The sounds were presented at a loudness of 60 dB, and the last 20 ms of the sounds were faded out in Audacity (version 2.4.2). In Experiments 1 and 3, an additional 41.5 Hz modulation was applied to the sounds to measure the ASSR. The startle probe, used in Experiments 1 and 3, was a 50 ms white noise burst presented at 95 dB.

### Procedure

Participants were instructed to abstain from eating and to drink only water for one hour prior to the experiment. They were also asked not to use any perfume or deodorant on the day of the experiment. Upon arrival, participants were briefed on the experimental procedure and completed questionnaires while electrodes were placed. The experiment began with an odor selection phase, where all odorants were presented twice and rated for intensity and pleasantness. Subsequently, all sounds, including the startle probe (in Experiments 1 and 3 only), were presented once and rated on the same scales. Based on the odor ratings, one of three predetermined pleasant odorants, i.e. Isoamyl acetate (smells like banana), Orange oil, or Lemon oil was selected as the US+appetitive stimulus. Butyric acid (smells like vomit) was used as the US+aversive stimulus, and pure mineral oil solution served as the US-. The odorants were randomly paired with the CS for the duration of the experiment.

Trials were pseudo-randomized to ensure no more than two consecutive trials of the same condition (CS+appetitive, CS+aversive, CS-) occurred. The 300 trials (100 per condition) were equally divided among 10 blocks with intervening breaks. Every second block was followed by a break during which participants filled out questionnaires, while alternate breaks allowed participants to briefly remove the nose pieces and drink water. Experiment 4 consisted of only one block with 10 trials per condition.

At the end of the experiment, contingency awareness was assessed through a semi-structured interview with hierarchical contingency questions and a contingency awareness questionnaire following the approach of Manns et al. (2001; see Supplementary material 1).

### Data acquisition and preprocessing

#### Behavioral measures

Participants rated pleasantness (−50 to 50) and intensity (0 to 100) of the odors after each trial and the CS sounds after each block on a VAS. Furthermore, the German translations of the following questionnaires (implemented via formr; Arslan et al., 2020) were used to assess personality traits: State-Trait Anxiety Inventory, Positive and Negative Affect Schedule, Stanford Sleepiness Scale, Intolerance of Uncertainty Scale, Arnett Inventory of Sensation Seeking, NEO Five-Factor Inventory, Behavioral Inhibition System and Behavioral Approach System. The questionnaire results can be found in Supplementary material 2.

#### Peripheral physiology

Peripheral physiological signals were recorded from AUX channels of the EEG amplifier (actiCHamp, Brain Products GmbH). We used BIP2AUX adapters with a gain of 100, and placed the ground for the physiological channels on the lower right side of the forehead (Figure 2). Data were recorded at a sampling rate of 1000 Hz and filtered offline with a finite impulse response filter using the Hamming window method (mne.filter.filter_data function in MNE-Python; version 1.3.1; Gramfort et al., 2013). MNE-Python was used for data preprocessing of all physiological measures and EEG.

**Figure 2.**
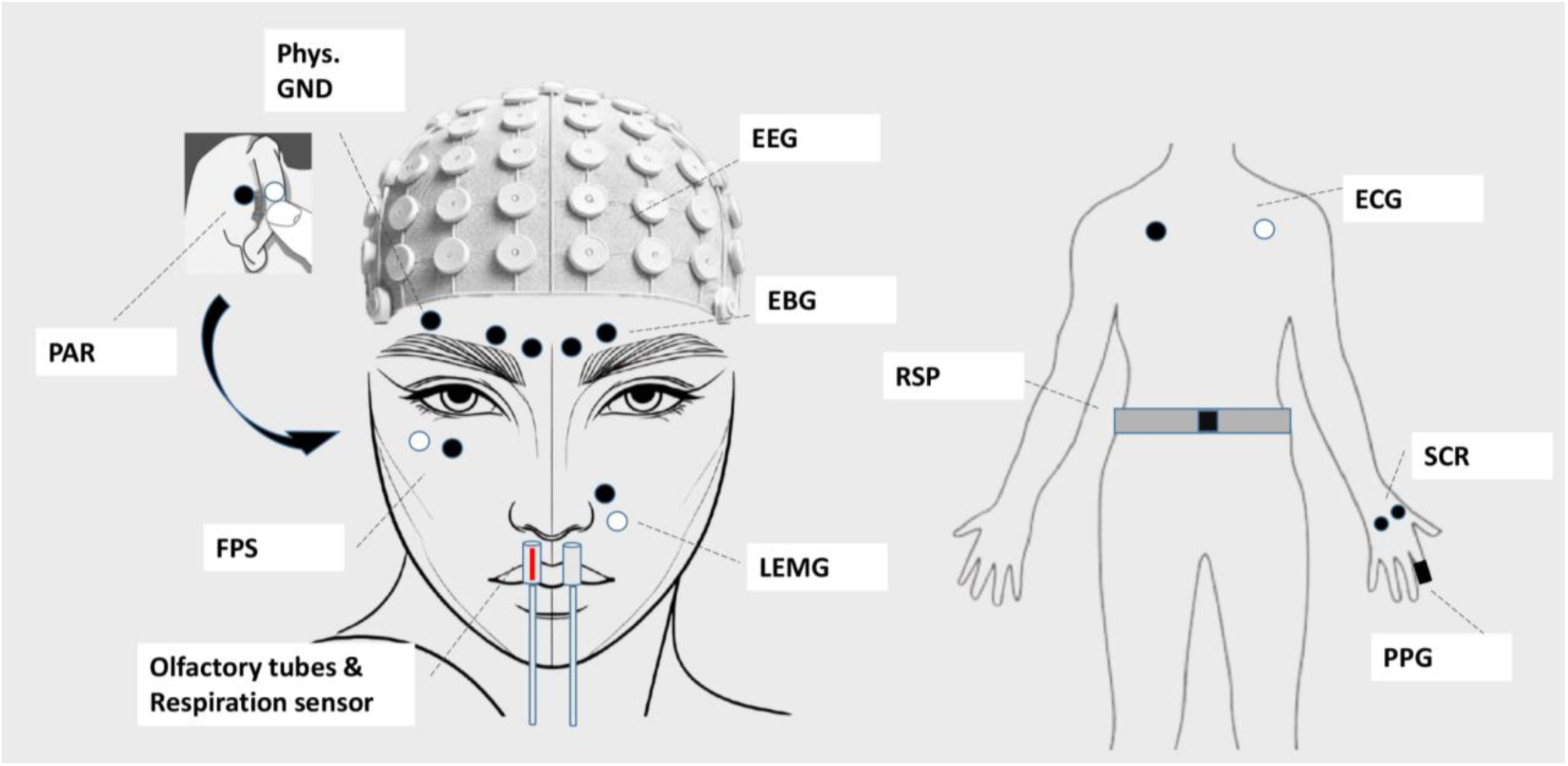
Placement of electrodes. For measures that were recorded with a bipolar montage (FPS, PAR, LEMG, and ECG), the black dot depicts the positive electrode and white dot the reference electrode. Phys. GND: physiological ground, PAR: postauricular reflex, FPS: fear-potentiated startle, EEG: electroencephalography, EBG: electrobulbogram, LEMG: levator labii superioris electromyography, RSP: respiration, ECG: electrocardiography, SCR: skin conductance response, PPG: photoplethysmography.

#### Skin conductance response

SCRs were measured using the GSR module (Brain Products GmbH) with electrodes on the lower thenar and hypothenar of the left hand. The signal was transformed online from microvolt to microsiemens using a gradient of 25. A 0.05 Hz high-pass- and a 5 Hz low-pass filter were applied to the raw signal before the data was epoched into [−2000 – 9500 ms] bins in Experiment 1 and 3, [−2000 – 7000 ms] bins in Experiment 2, and [−2000 – 8000 ms] bins in Experiment 4. The signal was then inspected manually for movement and noise artifacts, and trials containing artifacts were removed. The baseline period was set to the 2 s before the stimulus onset. The data were then baseline corrected following the approach of Kuhn et al. (2022), where the average of the baseline was subtracted from the peak, i.e. maximum value after CS onset, and the resulting difference was used for statistical analyses. If the resulting difference was a negative number, it was set to 0. Due to malfunction of the GSR module, 9 participants in experiment 3, and 7 participants in experiment 4 were removed from further analyses which led to a strong decrease in sample size in these two experiments. We still report the statistics, however urge for a limited interpretation of them.

#### Heart rate

ECG was recorded using a BIP2AUX adapter with electrodes on the left and right intersection of the midclavicular and subclavicular line (Figure 2). The signal was filtered with a 0.5 Hz high-pass filter, a 30 Hz low-pass filter, and a 50 Hz notch filter to improve R-peak detection. The data were epoched into [−4000 – 19,000 ms] bins in Experiment 1 and 3, [−4000 – 16,000 ms] bins in Experiment 2, and [−4000 – 11,000 ms] bins in Experiment 4, where 0 represented the CS onset. Automatic R-peak detection with manual correction was performed with HRVTool (Vollmer, 2019). R-peaks were then transformed to RR intervals, and then converted to instantaneous beats per minute (BPM). After that, the epochs for analysis were cropped into [−1000 – 5000 ms] bins. The reason for initially having larger bins was to account for long RR intervals at the epochs’ edges, and to inspect responses to the US. BPM values above 220 or below 50 were removed from the data. Then, the epoch was divided into 500 ms bins (i.e. 12 bins with an epoch length of 6 s), and for each bin we checked whether one or multiple BPM values were present. If only less than half of the bins contained a BPM value, the trial was rejected. Otherwise, the signal was interpolated to create an epoch containing one BPM value per bin, and offering a temporal development of the heart rate. The resulting epochs were then baseline-corrected per participant by calculating the average baseline signal for each CS condition, and subtracting it from the signal. We then analyzed the average heart rate change over the first deceleration [0 – 1000 ms], first acceleration [1000 – 2000 ms], second deceleration [2000 – 5000 ms], and the entire CS duration [0 – 5000 ms] after CS onset.

#### Pulse wave amplitude

Photoplethysmography (PPG) was measured using a PPG module (FpSens A1; MKS) on the left index finger (Figure 2). The data were filtered with a 0.5 Hz high-pass filter and a 15 Hz low-pass filter. The data were epoched in the same way as the heart rate (see above), where originally larger epochs were created than the ones used for the analysis. The systolic peak and the preceding base were detected with HRVTool (Vollmer, 2019). The pulse wave amplitude (PWA) was then calculated by subtracting the base from the peak amplitude. Values above or below 3 SD of the mean were removed from the data. Then, similar to the heart rate preprocessing, we created 500 ms bins, and interpolated the PWA values in each epoch, rejecting trials if less than half of the bins contained PWA values. Participants, where more than half of the trials were rejected, were excluded from analyses. Similar to the heart rate baseline correction, the PWA was corrected by calculating the average of each CS condition and subtracting it from the signal. The PWA was analyzed over a period of [0 – 5000 ms] after CS onset.

#### Respiration

Respiration was measured using a respiration belt (Brain Products GmbH). The respiration belt was placed around the waist between the lowest rib and the navel (Figure 2). A 5 Hz low-pass filter was applied to the raw signal, and it was manually inspected for artifacts after epoching the data into [−2000 – 6000 ms] bins, where 0 represents the CS onset. Each epoch was baseline corrected, by subtracting the average of the baseline. The respiration amplitude was calculated by averaging over the entire CS duration [0 – 5000 ms]. The respiration rate was calculated using Neurokit2 (version 0.2.10), where the inter-breath-intervals were calculated for the entire epoch. The resulting data was then baseline corrected and averaged over the entire CS duration [0 – 5000 ms].

#### Fear-potentiated startle

The FPS was measured using a BIP2AUX adapter, placing one electrode below the pupil of the right eye and the other to the right of it (Figure 2). A 28 Hz high-pass filter, and a 499 Hz low-pass filter were applied to the raw signal, after which data were epoched into [−500 – 1500 ms] bins where 0 represents the startle onset. The signal was then rectified, and an additional 40 Hz low-pass filter was applied. Trials were manually inspected for artifacts. The epochs were then cropped to [−50 – 150 ms] intervals. The peaks were automatically extracted by taking the highest value in an epoch, and these peaks were then transformed to T-scores within participants and conditions for analysis (Blumenthal et al., 2005).

#### Postauricular reflex

The PAR was measured using a BIP2AUX adapter, placing one electrode on the postauricular muscle behind the right ear, and the other one on the pinnacle as close as possible to the first electrode (Figure 2). The data were then preprocessed in the same way as the FPS, resulting in epochs with a time interval of [−50 – 150 ms]. The peaks were automatically extracted by taking the highest value in an epoch, and the peak-to-baseline difference was calculated within each trial per subject and then averaged for analysis.

#### Levator EMG

The levator labii superioris EMG was measured using a BIP2AUX adapter, placing one electrode on the middle of the levator labii superioris alaeque nasi on the left side of the nose, and the other electrode below it (Figure 2). A 499 Hz low-pass filter was applied to the raw signal, and all trials with a peak-to-peak signal amplitude higher than 30 mV were rejected automatically. This liberal threshold was based on the suggestion by Raez et al. (2006) that an EMG signal has a range of 0 – 10 mV (+5 or −5). The remaining trials were then rectified, and a 40 Hz low-pass filter was applied. Following the recommendations of Rutkowska et al. (2023), the muscle activity of the levator muscle was then baseline-corrected by dividing by the average baseline amplitude. The response was measured as the mean amplitude in the time window [0 – 5000 ms] after CS onset, except in Experiment 4 where the time window was [0 – 4500 s].

#### Corrugator EMG

The muscle activity at the left corrugator supercilii was extracted from the EEG electrodes that were placed there initially to measure the electrobulbogram (EBG; Figure 2). The electrode closest to the midline of the face above the left eyebrow was used for the signal, and the electrode further away from the midline above the left eyebrow was used as reference. After recording, the signal was re-referenced to this reference. The signal was then filtered with a 20 Hz high-pass filter and a 499 Hz low-pass filter. Similar to the levator EMG, we followed the recommendations of Rutkowska et al. (2023) for analysis. The signal of the corrugator EMG was baseline-corrected by dividing by the average baseline amplitude. Responses are the mean muscle amplitude in the time window [0 – 5000 ms] after the CS onset.

#### EEG

We recorded EEG with the ActiCHamp 64 channel system (Brain Products GmbH). Electrodes were placed according to the 10-20 system with Cz channel as the online reference and Fpz as the ground electrode. We placed four electrodes (PO3, PO4, PO7, and PO8) above the eyebrows for an EBG (Iravani et al., 2020). Impedances were maintained below 25 kOhm. The sampling rate was 1000 Hz.

Each recording was filtered offline by applying 0.01 Hz high-pass and 45 Hz low-pass filters. Data were re-referenced to average reference and epoched into [−2000 – 15,000 ms] bins in Experiment 1 and 3, and into [−2000 – 12,000 ms] bins in Experiment 2, with 0 representing the CS onset. Epochs containing high-amplitude artifacts that would distort the Independent Component Analysis (ICA) too much were visually identified and discarded. On average 4.46 % of all trials were excluded in Experiment 1, 4.28 % in Experiment 2, and 7.44% in Experiment 3. Noisy channels were excluded to later be interpolated, and then, an ICA was performed using the Picard algorithm. As a preliminary step for improving ICA decomposition, a 1 Hz high-pass filter was applied. The results of the ICA on the 1 Hz filtered data were then imported to the data filtered with the 0.01 Hz high-pass filter. Components clearly related to blinks, eye movements and high amplitude muscle activity were removed. Then, the excluded channels were interpolated (interpolate_bads function in MNE-Python), and reintegrated in the dataset. No EEG was recorded in Experiment 4.

#### Event-related potential

We report the difference in amplitude at the midline electrodes (Fz, Cz, Pz) at N1 [80 – 120 ms], P2 [180 – 220 ms], the LPP [500 – 700 ms], and over the entire CS duration without early components [1000 – 5000 ms], and the stimulus-preceding negativity (SPN) before the US onset [−1000 – 0 ms]. For this analysis the EEG signal was re-referenced to the average of both mastoid electrodes (TP9 and TP10).

#### Time-frequency analysis

We analyzed the frequency domain as well (1 – 45 Hz, Morlet wavelet transform), where visual inspection of the time-frequency representations (averaged across conditions) showed higher activity compared to baseline in the alpha band (8-12 Hz) at parietal electrodes (P1, Pz, P2), and suppression in the lower beta band (12-16 Hz) at occipital electrodes (O1, Oz, O2) during the CS presentation. We analyzed the time window [1000 – 5000 ms] after the CS onset in order to not include early ERP components of the CS response. Baseline normalization of the data was performed by subtracting the average baseline values [−1500 – −100 ms], and then divided by the average baseline.

#### Auditory steady-state response (ASSR)

To analyze the ASSR to the modulated frequency of the CS in Experiment 1 and 3, we followed the recommendations of Cohen and Gulbinaite (2017), where the rhythmic entrainment source separation (RESS) method was applied to optimize the signal-to-noise ratio (SNR) of the modulation frequency (41.5 Hz). After transforming the epoched data [1000 – 5000 ms] with the RESS method, the power spectrum was computed (0.5 – 45 Hz, Welch’s method). The power spectrum was then transformed to a SNR spectrum based on neighboring frequencies. These SNR values were used to statistically analyze the difference between CS conditions.

#### Statistical analysis

We used the Pingouin package (version 0.5.3) to conduct ANOVAs. In all measures, except EEG, but including ASSR, we tested the difference between conditions with a one-way RM ANOVA (with three levels of factor condition: appetitive CS+, aversive CS+, and CS-). For analyses of the ERP and time-frequency representations, we tested statistical differences by also taking the electrode position (Pz, Cz, and Fz) into account, resulting in a two-way RM ANOVA (condition x electrode). We also used permutation-based clustering in EEG (mne.stats.spatio_temporal_cluster_1samp_test in MNE-Python; 1024 permutations; t-threshold was determined with the scipy.stats.rv_continuous.ppf function) to identify significant clusters in time, space or frequency, and compared each of the conditions in a pair-wise manner with each other.

The assumption of sphericity was violated in some experiments, and in that case greenhouse-geisser corrected *p*-values are reported. Also, due to the exploratory nature of our study, the significance threshold was set at *p* <. 01. Whenever post-hoc comparisons are performed, we report the Benjamin-Hochberg FDR-corrected *p*-values for the *t*-tests.

In Experiments 1 and 2, not all participants were contingency aware. We do not report separate statistics for unaware or aware subjects in these experiments because the resulting sample sizes would be too small.

## Results

### Behavioral measures

#### CS ratings

Participants were asked to rate the CS (sounds) after every block, leading up to a total of 10 ratings of which the averages are reported here. Behavioral CS pleasantness ratings were not different between conditions in any of the four experiments (Figure 3). The ratings of the CS remained neutral (i.e. around a value of 0) throughout the experiment, and most importantly, they did not differ between the appetitive or aversive CS+, and no difference compared to the CS- was found either.

**Figure 3.**
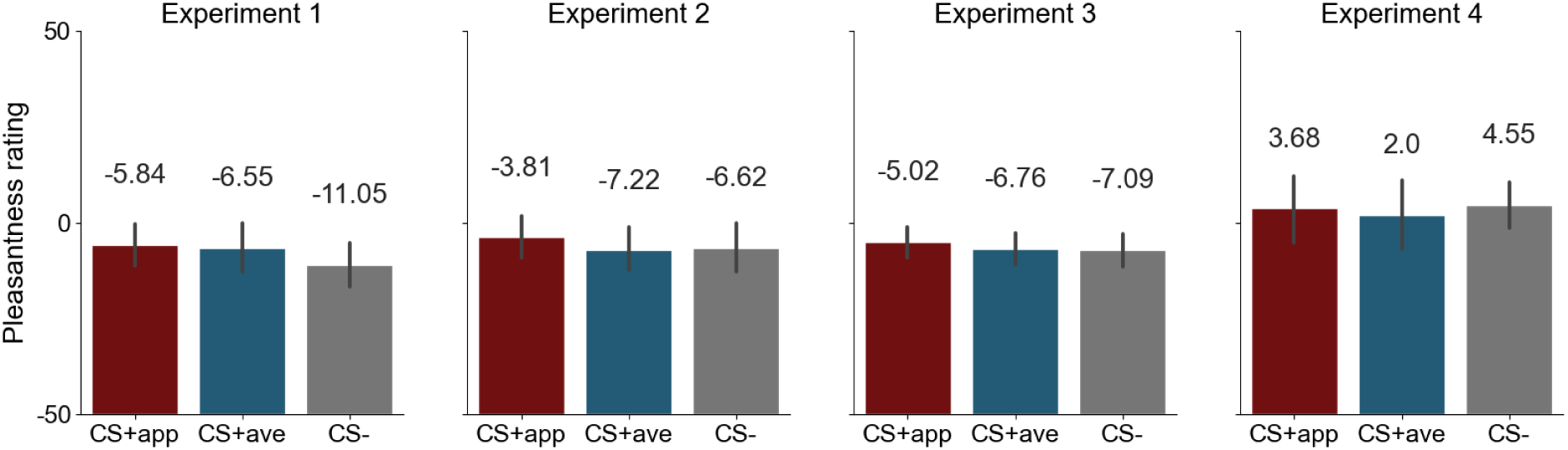
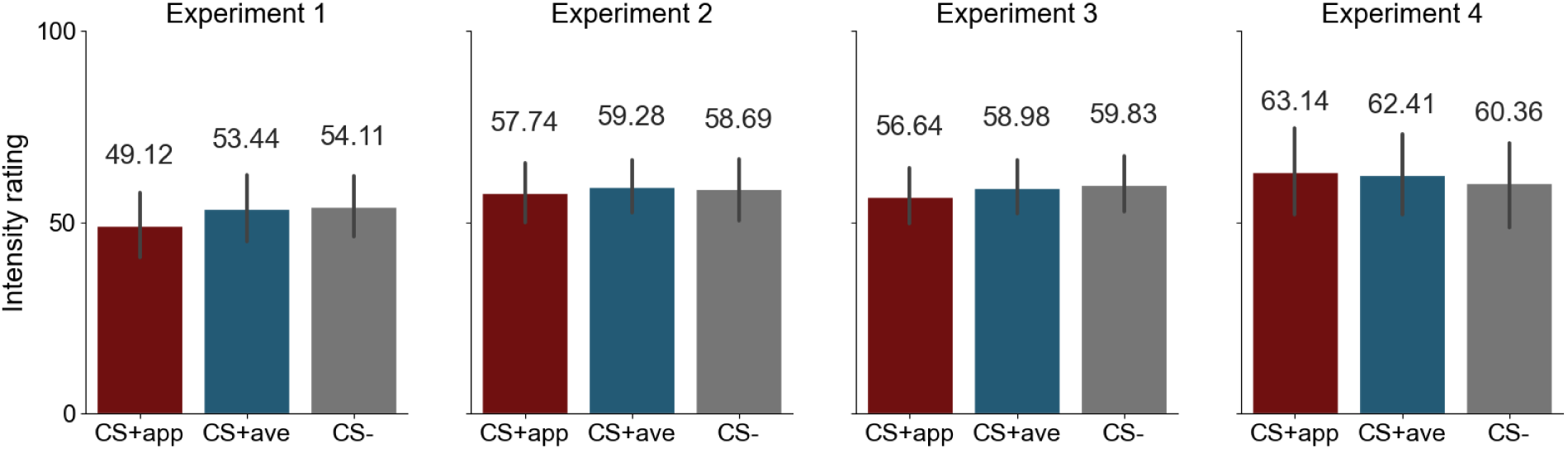
Pleasantness and intensity ratings of the CS+appetitive, CS+aversive, and CS-in each experiment. Error bars represent 95% CI.

When looking at Experiment 1, there was a significant difference between conditions in the intensity ratings of the CS (see Figure 3 and Table 2). Post-hoc comparisons showed, however, that the difference between the appetitive CS+ and the aversive CS+, and between the appetitive CS+ and CS-did not attain the significance level (both corrected *p* = .03), and no difference was observed between the aversive CS+ and CS-(corrected *p* = .55). There was also a significant effect in Experiment 3, however post-hoc comparisons revealed that there was no significant difference between intensity ratings in the three conditions (all corrected *p* > .015). In Experiment 2 and 4, the ANOVA indicated no difference between conditions.

**Table 2.** Results from the ANOVA for the pleasantness and intensity ratings of the CS and US in the four experiments.

| Measure | Experiment | CS |  |  | US |  |  |
| --- | --- | --- | --- | --- | --- | --- | --- |
| | | *F* (*df*) | *p* | $\eta_p^2$ | *F* (*df*) | *p* | $\eta_p^2$ |
| Pleasantness | 1 (*n*=19) | 4.64 (2, 36) | .016 | 0.21 | 53.12 (2, 36) | < .001 | 0.75 |
|  | 2 (*n*=20) | 3.30 (2, 38) | .048 | 0.15 | 51.82 (2, 38) | < .001 | 0.73 |
|  | 3 (*n*=32) | 1.04 (2, 62) | .36 | 0.03 | 146.16 (2, 62) | < .001 | 0.83 |
|  | 4 (*n*=22) | 0.11 (2, 42) | .90 | 0.01 | 132.33 (2, 42) | < .001 | 0.86 |
| Intensity | 1 (*n*=19) | 5.94 (2, 36) | .006 | 0.25 | 60.00 (2, 36) | < .001 | 0.77 |
|  | 2 (*n*=20) | 1.03 (2, 38) | .37 | 0.05 | 47.31 (2, 38) | < .001 | 0.71 |
|  | 3 (*n*=32) | 5.10 (2, 62) | .009 | 0.14 | 63.10 (2, 62) | < .001 | 0.67 |
|  | 4 (*n*=22) | 0.34 (2, 42) | .66 | 0.02 | 42.65 (2, 42) | < .001 | 0.67 |

#### US ratings

To investigate whether the valence or intensity of the US (odors) was insufficient for conditioning, we analyzed US pleasantness and intensity, which participants rated in every trial. The US pleasantness ratings (Figure 4) were highly significantly different between conditions in all experiments (Table 2). All post-hoc comparisons using *t-*test indicated corrected *p* < .001. Similar to the pleasantness ratings, the intensity ratings also indicated highly significant differences between conditions in all experiments. As expected, the intensity did not differ between appetitive and aversive US+ in any experiment, with all corrected *p* > .1. However, both the appetitive and aversive US+ were more intense than the control stimulus (US-), where all tests yielded corrected *p* < .001.

**Figure 4.**
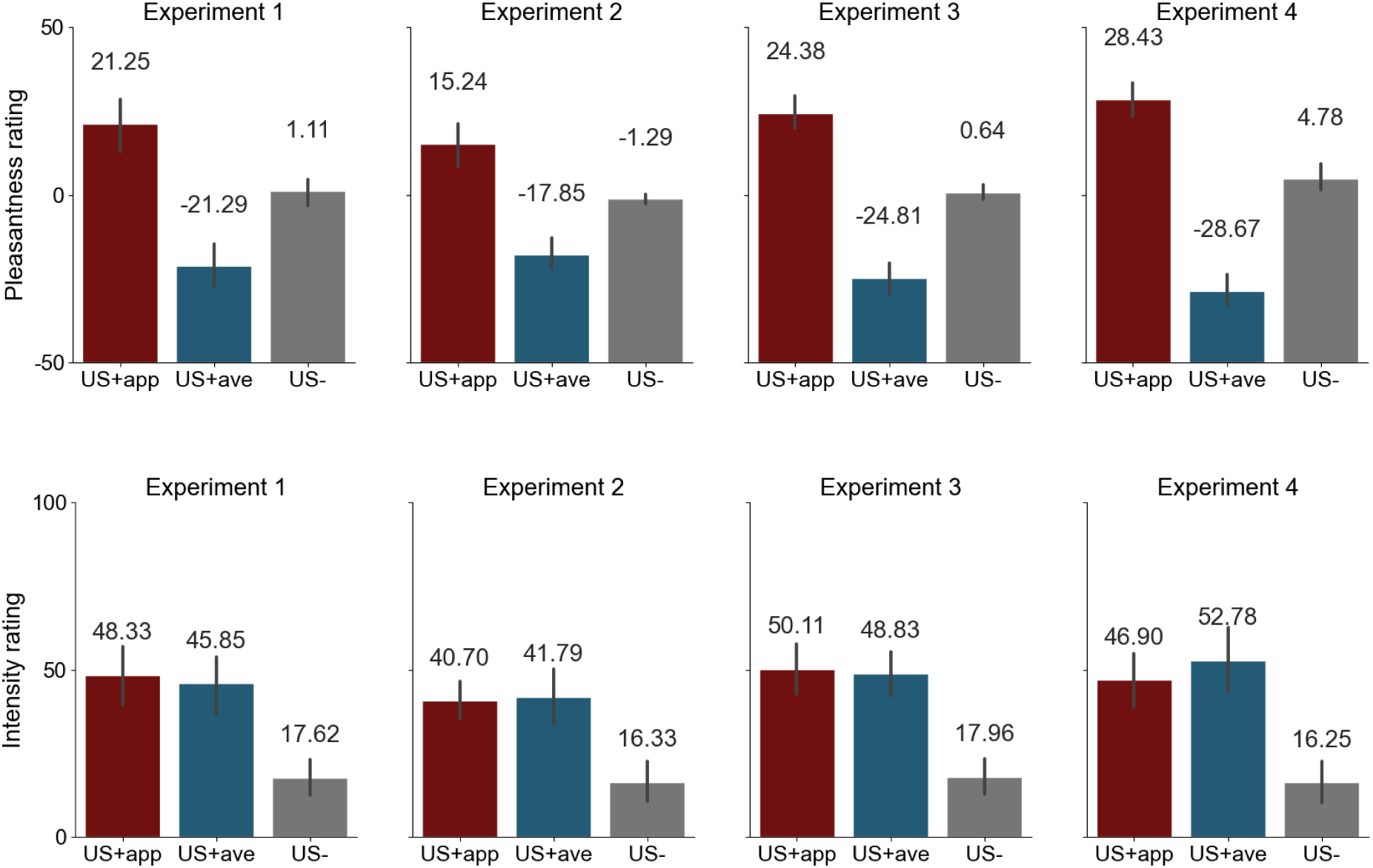
Pleasantness and intensity ratings of the US+appetitive, US+aversive, and US-in each experiment. Error bars represent 95% CI.

### Peripheral physiology

#### Skin conductance response

SCRs to the CS were not different between conditions in any of the experiments (see Table 3).

**Table 3.** Results from the ANOVA for each peripheral physiological measure in the four experiments.

| Measure | Experiment | $F(df)$ | $p$ | $\eta_p^2$ | Measure | Experiment | $F(df)$ | $p$ | $\eta_p^2$ |
| --- | --- | --- | --- | --- | --- | --- | --- | --- | --- |
| SCR | 1 ($n=17$) | 0.12 (2, 32) | .89 | 0.01 | HR (1st dec.) | 1 ($n=18$) | 0.62 (2, 34) | .54 | 0.04 |
|  | 2 (n=20) | 1.44 (2, 38) | .25 | 0.07 |  | 2 (n=20) | 0.78 (2, 34) | .44 | 0.04 |
|  | 3 (n=12) | 0.84 (2, 22) | .45 | 0.07 |  | 3 (n=30) | 0.69 (2, 58) | .50 | 0.02 |
|  | 4 (n=7) | 1.95 (2, 12) | .18 | 0.25 |  | 4 (n=22) | 2.17 (2, 40) | .13 | 0.10 |
| HR (1st acc.) | 1 (n=18) | 1.91 (2, 34) | .16 | 0.10 | HR (2nd dec.) | 1 (n=18) | 0.85 (2, 34) | .41 | 0.05 |
|  | 2 (n=20) | 1.78 (2, 34) | .20 | 0.09 |  | 2 (n=20) | 3.23 (2, 34) | .05 | 0.16 |
|  | 3 (n=30) | 0.14 (2, 58) | .87 | 0.01 |  | 3 (n=30) | 0.73 (2, 58) | .48 | 0.02 |
|  | 4 (n=21) | 0.46 (2, 40) | .63 | 0.02 |  | 4 (n=21) | 0.12 (2, 40) | .89 | 0.01 |
| HR (entire CS) | 1 (n=18) | 0.83 (2, 34) | .41 | 0.05 | PWA (entire CS) | 1 (n=16) | 1.82 (2, 30) | .18 | 0.11 |
|  | 2 (n=20) | 3.31 (2, 34) | .05 | 0.16 |  | 2 (n=10) | 0.95 (2, 18) | .41 | 0.10 |
|  | 3 (n=30) | 0.45 (2, 58) | .64 | 0.02 |  | 3 (n=31) | 2.25 (2, 60) | .11 | 0.07 |
|  | 4 (n=21) | 0.13 (2, 40) | .87 | 0.01 |  | 4 (n=21) | 0.59 (2, 40) | .56 | 0.03 |
| RSP amplitude (entire CS) | 1 (n=19) | 1.02 (2, 36) | .37 | 0.05 | RSP rate (entire CS) | 1 (n=19) | 0.51 (2, 36) | .61 | 0.03 |
|  | 2 (n=20) | 0.73 (2, 38) | .49 | 0.04 |  | 2 (n=20) | 0.26 (2, 38) | .77 | 0.01 |
|  | 3 (n=32) | 1.28 (2, 62) | .29 | 0.04 |  | 3 (n=31) | 0.64 (2, 60) | .49 | 0.02 |
|  | 4 (n=22) | 0.04 (2, 42) | .96 | < 0.01 |  | 4 (n=21) | 2.25 (2, 40) | .12 | 0.10 |
| FPS | 1 (n=18) | 1.47 (2, 34) | .24 | 0.08 | PAR | 1 (n=19) | 0.35 (2, 36) | .63 | 0.02 |
|  | - | - | - | - |  | - | - | - | - |
|  | 3 (n=31) | 7.03 (2, 60) | .002 | 0.19 |  | 3 (n=32) | 0.90 (2, 62) | .40 | 0.03 |
|  | - | - | - | - |  | - | - | - | - |
| LEMG | 1 (n=19) | 2.51 (2, 36) | .10 | 0.12 | CEMG | 1 (n=19) | 1.99 (2, 36) | .15 | 0.10 |
|  | 2 (n=20) | 0.02 (2, 38) | .98 | < 0.01 |  | 2 (n=20) | 2.51 (2, 38) | .09 | 0.12 |
|  | 3 (n=32) | 0.50 (2, 62) | .61 | 0.02 |  | 3 (n=32) | 0.21 (2, 62) | .81 | 0.01 |
|  | 4 (n=22) | 3.40 (2, 42) | .04 | 0.14 |  | 4 (n=22) | - | - | - |

*Note:* SCR: skin conductance response, HR: heart rate, dec.: deceleration, acc.: acceleration, PWA: pulse wave amplitude, RSP: respiration, FPS: fear-potentiated startle, PAR: postauricular reflex, LEMG: levator EMG, CEMG: corrugator EMG.

#### Heart rate

There were no significant differences between conditions during the first deceleration period, acceleration period, second deceleration period, or the entire CS duration in any experiment (Figure 5 and Table 3).

**Figure 5.**
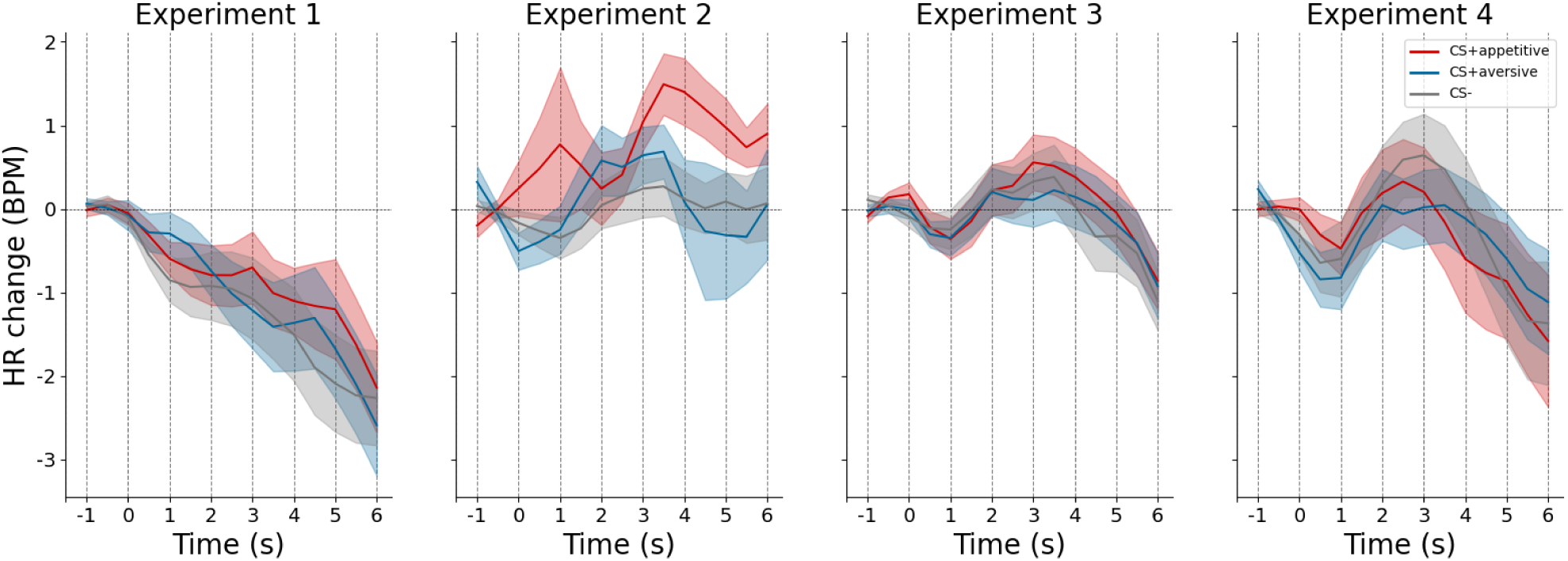
Heart rate response to the CS in all four experiments. Shaded areas represent the SEM.

#### Pulse wave amplitude

The average amplitude over the entire CS duration was compared between conditions, but showed no significant differences in any experiment (Figure 6 and Table 3).

**Figure 6.**
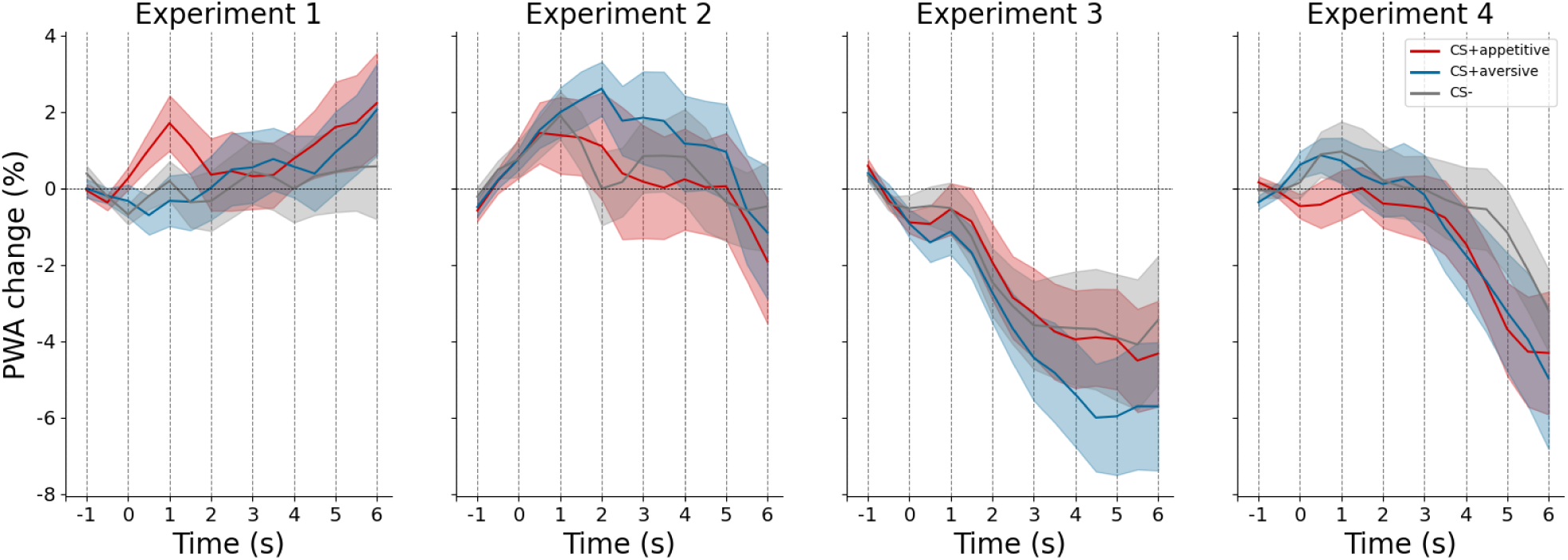
Percentual change of the pulse wave amplitude after the CS compared to baseline in all four experiments. Shaded areas represent the SEM.

#### Respiration

The respiration amplitude and respiration rate averaged over the entire CS duration showed no differences between CS conditions in any of the four experiments (Table 3).

#### Fear-potentiated startle

The FPS response was not different between conditions in Experiment 1 (Figure 7 and Table 3). However in Experiment 3, the FPS following the aversive CS+ elicited a significantly stronger response than the startle after the appetitive CS+ (corrected *p* = .01, *d* = 1.05) and after CS-(corrected *p* = .01, *d* = 0.87). There was no difference between the appetitive CS+ and CS-(corrected *p* = .35, *d* = 0.28). No startle probes were presented in Experiments 2 and 4.

**Figure 7.**
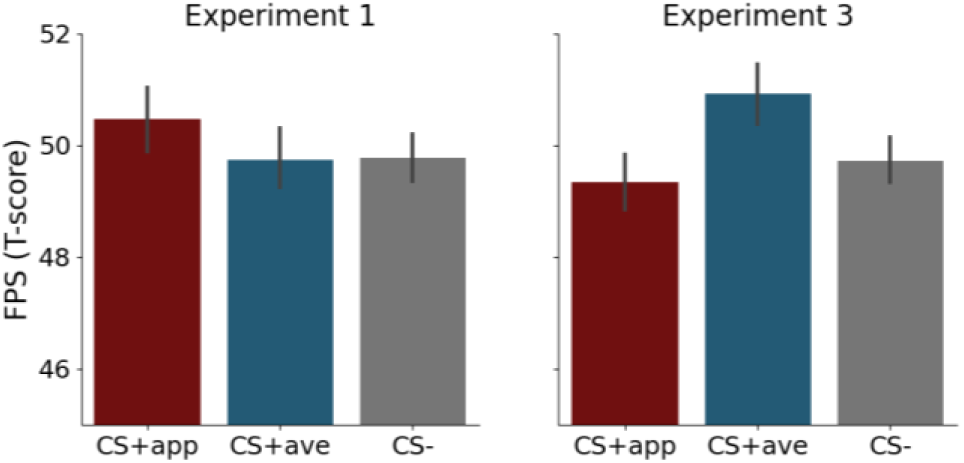
Fear-potentiated startle response in each experiment to the CS+appetitive, CS+aversive, and CS-. Error bars represent 95% CI.

#### Postauricular reflex

Contrary to the FPS, the reflex response of the post-auricular muscle was not modulated by the CS in Experiment 1 and 3 (Table 3).

#### Levator EMG

In none of the experiments the levator muscle activity differed between the CS (Table 3).

#### Corrugator EMG

Similar to the levator EMG, the CS did not evoke a different response in the corrugator muscle in any of the experiments. No corrugator EMG was recorded in Experiment 4 (Table 3).

### EEG

#### ERP

For the N1, P2, in Experiment 1, 2, and 3, there was no main effect of condition, but a significant main effect of electrode, yet no interaction effect of condition and electrode (Table 4 and Figure 8). The main effect of electrode for the LPP was only significant in Experiments 2 and 3. For the entire CS duration and SPN, there were no significant effects in any of the experiments.

**Figure 8.**
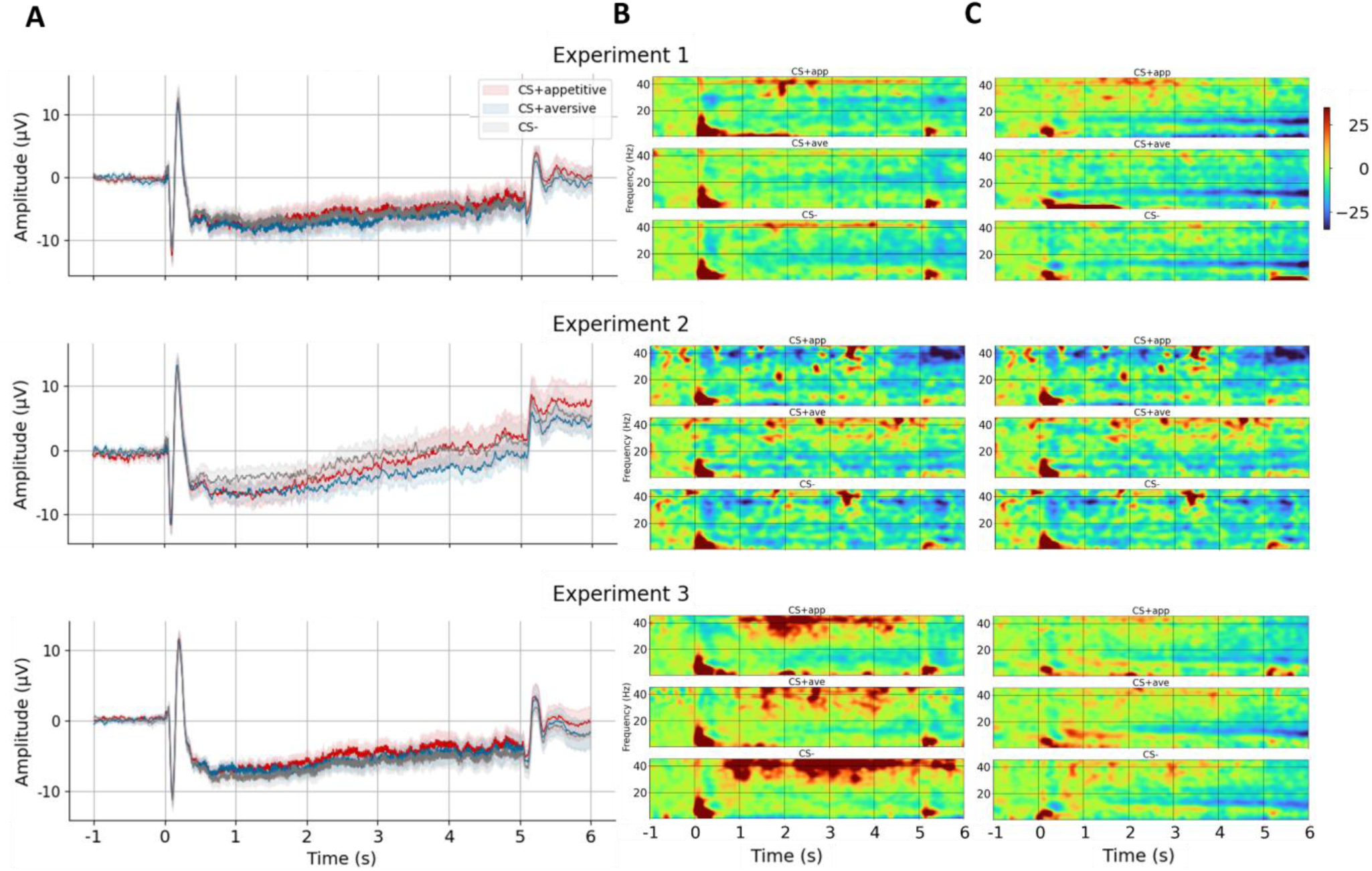
Event-related potentials (ERP) and time-frequency responses (TFR) in Experiments 1, 2, and 3. **A.** ERP to the CS at the Cz electrode. Shaded areas represent the SEM. **B**. Average TFR of the CS response at electrodes Cz, C1, and C2. **C.** Average TFR of the CS response at electrodes Oz, O1, and O2. The TFR colormap depicts the percentual power change to baseline.

**Table 4.** Results from the ANOVA for the ERP components in Experiment 1, 2, and 3.

| Measure | Experiment | Condition |  |  | Electrode |  |  | Condition x electrode |  |  |
| --- | --- | --- | --- | --- | --- | --- | --- | --- | --- | --- |
| | | *F* (df) | *p* | $\eta_p^2$ | *F* (df) | *p* | $\eta_p^2$ | *F* (df) | *p* | $\eta_p^2$ |
| N1 | 1 (n=18) | 0.35 (2, 34) | .67 | 0.02 | 14.49 (2, 34) | < .001 | 0.46 | 1.20 (4, 68) | .30 | 0.07 |
|  | 2 (n=18) | 1.29 (2, 34) | .28 | 0.057 | 15.56 (2, 34) | < .001 | 0.48 | 0.48 (4, 68) | .56 | 0.03 |
|  | 3 (n=29) | 0.51 (2, 56) | .59 | 0.02 | 18.35 (2, 56) | < .001 | 0.40 | 0.89 (4, 112) | .41 | 0.03 |
| P2 | 1 (n=18) | 0.52 (2, 34) | .57 | 0.03 | 24.37 (2, 34) | < .001 | 0.59 | 1.36 (4, 68) | .27 | 0.07 |
|  | 2 (n=18) | 0.03 (2, 34) | .97 | < 0.01 | 36.18 (2, 34) | < .001 | 0.68 | 0.62 (4, 68) | .51 | 0.03 |
|  | 3 (n=29) | 0.93 (2, 56) | .40 | 0.03 | 47.27 (2, 56) | < .001 | 0.63 | 0.81 (4, 112) | .44 | 0.03 |
| LPP | 1 (n=18) | 1.16 (2, 34) | .32 | 0.06 | 4.25 (2, 34) | .03 | 0.20 | 2.96 (4, 68) | .08 | 0.15 |
|  | 2 (n=18) | 1.11 (2, 34) | .32 | 0.06 | 6.01 (2, 34) | .008 | 0.26 | 0.65 (4, 68) | .48 | 0.04 |
|  | 3<br>(*n*=29) | 1.66 (2, 56) | .20 | 0.06 | 38.21 (2, 56) | < .001 | 0.58 | 0.82 (4, 112) | .44 | 0.03 |
| Entire CS<br>(1000–5000 ms) | 1<br>(*n*=18) | 0.86 (2, 34) | .43 | 0.05 | 0.20 (2, 34) | .75 | 0.01 | 1.54 (4, 68) | .23 | 0.08 |
|  | 2<br>(*n*=18) | 1.37 (2, 34) | .27 | 0.07 | 2.44 (2, 34) | .11 | 0.13 | 0.46 (4, 68) | .60 | 0.03 |
|  | 3<br>(*n*=29) | 1.61 (2, 56) | .21 | 0.05 | 3.37 (2, 56) | .05 | 0.11 | 0.72 (4, 112) | .47 | 0.03 |
| SPN | 1<br>(*n*=18) | 0.24 (2, 34) | .67 | 0.01 | 0.60 (2, 34) | .53 | 0.03 | 0.62 (4, 68) | .46 | 0.04 |
|  | 2<br>(*n*=18) | 0.56 (2, 34) | .50 | 0.03 | 1.06 (2, 34) | .35 | 0.06 | 0.49 (4, 68) | .50 | 0.03 |
|  | 3<br>(*n*=29) | 0.10 (2, 56) | .90 | < 0.01 | 1.58 (2, 56) | .22 | 0.05 | 0.63 (4, 112) | .51 | 0.02 |

Note: LPP: late positive potential, SPN: stimulus-preceding negativity.

#### TFR

Statistical analyses of the alpha power at parietal electrodes revealed no significant differences between conditions in either Experiment 1, 2 or 3. The lower beta power over occipital electrodes also was not significantly different between conditions in any experiment (Table 5 and Figure 8).

**Table 5.** Results from the ANOVA for the EEG time-frequency representations in experiment 1, 2, and 3.

| Experiment | Parietal Alpha |  |  | Occipital Beta |  |  | ASSR |  |  |
| --- | --- | --- | --- | --- | --- | --- | --- | --- | --- |
| | *F* (*df*) | *p* | $\eta_p^2$ | *F* (*df*) | *p* | $\eta_p^2$ | *F* (*df*) | *p* | $\eta_p^2$ |
| 1 (*n*=18) | 0.67 (2, 34) | .52 | 0.04 | 1.13 (2, 34) | .33 | 0.06 | 1.14 (2, 34) | .33 | 0.06 |
| 2 (*n*=18) | 1.71 (2, 34) | .20 | 0.09 | 1.93 (2, 34) | .10 | 0.02 | 0.82 (2, 34) | .45 | 0.05 |
| 3 (*n*=29) | 1.83 (2, 56) | .17 | 0.06 | 0.88 (2, 56) | .42 | 0.03 | 0.32 (2, 56) | .69 | 0.01 |

*Note:* ASSR: auditory steady-state response.

Additionally, the EEG signal was analyzed using permutation-based spatio-temporal frequency clustering of all electrodes, where we looked at the differences between conditions over the entire epoch [0 – 11,000 ms]. No significant clusters could be identified in any of the experiments.

#### ASSR

No significant differences of auditory steady-state response (ASSR) were found between conditions in any of the three experiments (Table 5).

## Discussion

The goal of our study was to investigate the physiological differences between appetitive and aversive conditioning in humans. To this end, we compared responses to sounds paired with pleasant, unpleasant, and odorless odors. We employed a wide range of physiological measures to explore potential differences in the underlying mechanisms of appetitive and aversive conditioning. However, aside from the FPS response observed in Experiment 3, neither the subjective ratings nor the physiological data across the four experiments showed any evidence of conditioning.

A possible reason why no conditioning occurred could be the affective properties of the US. However, we could rule out an insufficiently strong or distinct olfactory US, because participants clearly differentiated between odors. Specifically, they rated pleasant odors as pleasant and unpleasant odors as unpleasant, and as more intense than the control in all experiments.

Furthermore, the conditioning process may have been hindered by certain properties of the CS and the experimental design. The frequency modulation of the sounds (used to elicit ASSR) could have complicated their perception. Additionally, startle stimuli inserted between the CS and US may have obscured the contingency between the two, while the long and variable trace between CS and US could have further diminished their associative strength. To address these concerns, we omitted startle probes and used non-frequency-modulated CS sounds in Experiment 2. However, the results of Experiments 1 and 2 were nearly identical, with no observable differences between CS+ and CS-conditions in any behavioral or physiological measure. Experiment 4 further validated that the lack of conditioning was not due to the long trace between CS and US. These findings suggest that the interference of the startle stimuli, CS modulation, and the long trace are unlikely to be the sole causes of failed conditioning in our experiments.

Moreover, the absence of conditioning could be attributed to the complexity of using both an appetitive and aversive CS in parallel. However, previous work showed successful conditioning when combining appetitive and aversive olfactory conditioning (Exner et al., 2021; Gottfried et al., 2002; Hermann et al., 2000). It should be noted though that some tested aversive and appetitive conditioning between subjects (Hermann et al., 2000) or had separate appetitive and aversive acquisition blocks (Exner et al., 2021).

When looking at other olfactory conditioning studies, it is not entirely unexpected that conventional measures like the SCR or heart rate showed no discriminable responses to the different CS. Especially in appetitive conditioning, the SCR seems to not show a significant response (Exner et al., 2021; Hermann et al., 2000, 2002; Stussi et al., 2018). Similarly, heart rate responses also seem to only differ in aversive conditioning (Exner et al., 2021; Flor et al., 2002; van den Bosch et al., 2015), but not always (Hermann et al., 2000, 2002). The absence of CRs in SCR or heart rate could be explained by their nature as measures of arousal.

Assuming that aversive conditions are generally more arousing than appetitive ones, this would explain why these measures only show differential responses in the aversive condition (Exner et al., 2021; Flor et al., 2002; Hermann et al., 2000, 2002; van den Bosch et al., 2015). Perhaps, unlike fear conditioning with electrical shocks as the US, the disgust caused by aversive odors is also not a strong activator of the sympathetic nervous system, which is implicated in the origin of SCR and partially explains heart rate responses.

Previous studies demonstrated an effect of slower and delayed breathing in expectation of an unpleasant odor (Resnik et al., 2011; Arzi et al., 2012). Resnik et al. (2011) found that respiration is modulated by a pure tone (250 ms) paired with an unpleasant odor. Respiratory CRs may be easier to achieve, and since Arzi et al. (2012) and Resnik et al. (2011) used relatively short auditory CS preceding the olfactory US, this short interval might be enough to adjust one’s breathing before the US, be it consciously or unconsciously. In Experiment 4 of our study, the CS onset was followed by the olfactory US after 4500 ms, and possibly due to this long interval, no conditioned respiration responses were found to the CS.

More valence-dependent measures, such as the PAR, FPS, and facial EMG, would be expected to display clearer differential responses across conditions. For example, Stussi et al. (2018) demonstrated an increase in PAR after pairing a visual stimulus with an appetitive odor but found no modulation of FPS in the appetitive condition. While no FPS modulation is typically expected in appetitive conditions, Hermann et al. (2000) also found no change in FPS in the aversive condition. However, in an aversive conditioning-only study, Hermann et al. (2002) and Flor et al. (2002) did observe FPS potentiation. Given that these two studies also found a significant difference in SCR, this suggests that their experimental design, rather than the measures themselves, was responsible for the successful conditioning. Hermann et al. (2000) also recorded EMG of the corrugator supercilii and zygomaticus, but found no difference between CS+ and CS-in contrast to Flor et al. (2002) who found an effect in the corrugator supercilii. Thus, while facial EMG of muscles such as the corrugator supercilii, zygomaticus, and levator labii have shown to be modulated by an odors’ valence (Delplanque et al., 2009; Kato & Yagi, 1994), they may not always establish as CRs. Similarly, EEG in appetitive and aversive olfactory conditioning also showed limited potential to differentiate between CS conditions.

Only the FPS response in Experiment 3 showed sensitivity to conditioning. However, this result should be interpreted cautiously due to the numerous other tests conducted in the study. Nevertheless, this finding aligns with previous research (Flor et al., 2002; Haan et al., 2018; Hermann et al., 2002; Kuhn et al., 2019), which reported differential aversive conditioning in the startle response, although startle potentiation is not always observed in similar conditions (Hermann et al., 2000). The question remains why no effect was found in Experiment 1. Since the only difference from Experiment 3, apart from a slightly larger sample size, was the contingency instruction. This suggests that awareness may be necessary for the effect to manifest itself. Some studies show that FPS modulation can only be observed in contingency-aware participants (Glenn et al., 2012), however, other studies have shown that FPS modulation can occur without contingency awareness (Hamm & Weike, 2005; Jovanovic et al., 2006; Sevenster et al., 2014). The startle response can be attenuated when attention is directed to another modality (Pastor et al., 2015; Schicatano & Blumenthal, 1998), which could explain why unaware participants might show weaker FPS potentiation, and why the effect only became apparent in Experiment 3.

Olfactory conditioning seems to be more fruitful during sleep as Canales-Johnson et al. (2020) found sniff differentiation for appetitive and aversive CS trials. Furthermore, Arzi et al. (2012) also found that participants could be conditioned during sleep, resulting in an inhalatory response to the CS (1000 ms pure tone) after waking. It seems contradictory that olfactory conditioning would work better during sleep. However, Cellini and Parma (2015) suggest that olfactory conditioning during sleep is easier because there is less sensory input to the cortex (Steriade, 2003), and the extra-thalamic olfactory inputs may have privileged access to the cortex. Furthermore, they cite Barnes and Wilson (2014) who have found that functional connectivity during this sleep stage increases between olfactory, limbic and neocortical areas. Further tests of the replicability of these findings are deemed necessary.

The nature of olfactory neural pathways can be another factor affecting olfactory conditioning. As previously noted, olfactory pathways do not require a thalamic relay for conscious processing (Shepherd, 2005). The extra-thalamic pathways leading to the amygdala and entorhinal cortex are expected to facilitate affective evaluation and memory formation during conditioning. However, this feature may hinder the association between an auditory CS and an olfactory US. This raises the question why olfactory conditioning with visual CS does not appear to face the same challenges (e.g. Exner et al., 2021; Hermann et al., 2002; Stussi et al., 2018). Unlike visual depictions of food or other odorous objects, there are few instances in real life where a sound automatically evokes an association with an odor. The lack of this hardwired connection between auditory and olfactory areas of the brain may explain our findings.

## Acknowledgments

This study was supported by the Else-Kröne-Fresenius Stiftung. We would further like to thank Simone Liebscher at the Schenke-Layland lab in Tübingen for her help in creating the olfactory stimuli.

## Data and code availability statement

All raw data and code of Experiment 1 can be retrieved at https://doi.org/10.12751/g-node.n972n6, of Experiment 2 at https://doi.org/10.12751/g-node.h5j17z, of Experiment 3 at https://doi.org/10.12751/g-node.j3nsmn, and of Experiment 4 at https://doi.org/10.12751/g-node.36ffx7. The behavioral data can be retrieved at https://osf.io/2hkz4/

## Funding statement

This project was funded by the Else-Kröne-Fresenius Stiftung.

## Conflict of interest disclosure

K.O. works for dsm-firmenich. The company had no role in the design of the studies and interpretation of results. The other authors have no conflicts of interest to disclose.

## CRediT statement

**NS Menger**: Conceptualization, Methodology, Software, Validation, Formal analysis, Investigation, Resources, Data curation, Writing - original draft, Writing - review & editing, Visualization

**YG Pavlov**: Conceptualization, Methodology, Software, Validation, Resources, Writing - original draft, Writing - review & editing, Supervision, Project administration

**B Kotchoubey**: Conceptualization, Methodology, Writing - original draft, Writing - review & editing, Supervision, Project administration, Funding acquisition

**K Ohla**: Conceptualization, Methodology, Writing - review & editing

## Supplementary material

**SM 1.** Contingency awareness interview and questionnaire

**SM 2.** Personality traits

## Notes

https://gin.g-node.org/nickmenger/Menger_et_al_2025_Experiment_1

https://gin.g-node.org/nickmenger/Menger_et_al_2025_Experiment_2

https://gin.g-node.org/nickmenger/Menger_et_al_2025_Experiment_3

https://gin.g-node.org/nickmenger/Menger_et_al_2025_Experiment_4

### Competing Interest Statement

K. Ohla works for dsm-firmenich. The company had no role in the design of the studies and interpretation of results. The other authors have no conflicts of interest to disclose.

