## Supplementary material for "Missing what’s right under your nose: failed appetitive and aversive audio-olfactory conditioning in humans": SM 1

**SM 1. Contingency awareness interview and questionnaire**

**Post-experiment interview:**

The interview questions were asked in the following order:

1. What did you perceive?
2. Was there anything else?
3. How many sounds did you hear?
4. How many odors did you smell?
5. Did you have the feeling that the odors were preceded by specific sounds?
6. If you had to guess which odor came after sound 1, then … which odor came after sound 2 then …, which odor came after sound 3, then …?
7. At which time point during the experiment did you notice the system/contingencies (rate on a scale of 1-100, where 1 is the beginning of the experiment and 100 is the end)?

If the participant did not get the contingencies, then they were explained to the participant.

**Post-experiment questionnaire:**

1. What did you perceive/feel during the experiment?

   *Participant could tick multiple boxes.*

   - Pleasant smells
   - Electric shocks
   - Unpleasant smells
   - Loud sounds
   - Pushing of the body
   - Various sounds
2. Did you notice something else?
   - Yes
   - No
3. The following questions are regarding the sounds and smells in the experiment. Please describe to what extent the statement applies to you. Please answer quickly but carefully.

   *Participant answers on a scale from "I don't agree at all" to "I fully agree"*

   I believe that the sound always **preceded** the smell.

   I believe that the sound always **succeeded** the smell.

   I believe that the smell always **preceded** the sound.

   I believe that the smell always **succeeded** the sound.

   I believe that the sounds and smells were always related temporally.

   I believe that the sounds and smells were sometimes related temporally.

   I believe that the sounds predicted when the smell came.

4. The following questions are regarding the sounds and smells in the experiment. Please describe to what extent the statements apply to you. Please answer quickly but carefully.

   *Participant answers on a scale from "completely unlikely" to "completely likely"*

   How high was the probability that the AAAAA-sound was followed by a pleasant smell?

   How high was the probability that the AAAAA-sound was followed by an unpleasant smell?

   How high was the probability that the AAAAA-sound was followed by no smell?

   How high was the probability that the EEEEE-sound was followed by a pleasant smell?

   How high was the probability that the EEEEE-sound was followed by an unpleasant smell?

   How high was the probability that the EEEEE-sound was followed by no smell?

   How high was the probability that the UUUUU-sound was followed by a pleasant smell?

   How high was the probability that the UUUUU-sound was followed by an unpleasant smell?

   How high was the probability that the UUUUU-sound was followed by no smell?
