## Supplementary material for "Missing what’s right under your nose: failed appetitive and aversive audio-olfactory conditioning in humans": SM 2

**SM 2. Personality traits**

**Table 1.** Mean (SD) personality trait scores in Experiments 1, 2, and 3.

| Experiment | STAI trait | IUS factor 1 | IUS factor 2 | AISS novelty | AISS intensity | NEO (N) | NEO (E) | NEO (O) | NEO (C) | NEO (A) | BIS/BAS factor 1 | BIS/BAS factor 2 | BIS/BAS factor 3 | BIS/BAS factor 4 |
|---|---|---|---|---|---|---|---|---|---|---|---|---|---|---|
| 1 (*n* = 18) | 35.9 (9.2) | 84.2 (28.8) | 83.3 (29.7) | 2.7 (0.3) | 2.2 (0.5) | 20.8 (6.5) | 26.2 (4.7) | 18.1 (3.1) | 30.9 (4.7) | 17.2 (2.2) | 12.0 (1.8) | 11.8 (2.0) | 16.4 (1.7) | 21.1 (3.1) |
| 2 (*n* = 19) | 39.2 (7.4) | 93.3 (25.0) | 84.6 (21.8) | 2.7 (0.4) | 2.2 (0.4) | 22.6 (6.1) | 26.9 (4.4) | 18.3 (3.2) | 30.1 (5.2) | 16.4 (2.4) | 12.2 (2.0) | 12.5 (2.7) | 17.3 (2.5) | 21.1 (4.3) |
| 3 (*n* = 32) | 37.0 (7.7) | 94.0 (27.4) | 92.4 (20.5) | 2.7 (0.5) | 2.3 (0.4) | 22.2 (5.2) | 26.8 (4.6) | 17.7 (3.0) | 30.6 (5.3) | 17.1 (1.7) | 11.9 (2.3) | 11.9 (2.2) | 16.4 (2.0) | 21.2 (3.3) |

*Note*: STAI: State-Trait Anxiety Inventory, IUS: Intolerance of Uncertainty Scale, AISS: Arnett Inventory of Sensation Seeking, NEO: NEO Five-Factor Inventory, BIS/BAS: Behavioral Inhibition System and Behavioral Approach System.

Before, during, and after Experiments 1, 2, and 3, participants rated their anxiety level (Figure 1). The STAI state anxiety level significantly changed over the duration of Experiment 3, $F(2, 62) = 15.44$, $p < .001$, η2p = 0.33 during the experiment. Post-hoc comparisons showed that the anxiety level increased between the start and middle of the experiment ($p < .001$), and decreased at the end of the experiment ($p = .01$). In experiments 1 and 2, there was no significant change of anxiety level during the experiment; in Experiment 1, $F(2, 34) = 2.83$, $p = .07$, η2p = 0.14, and Experiment 2, $F(2, 36) = 2.42$, $p = .10$, η2p = 0.12.

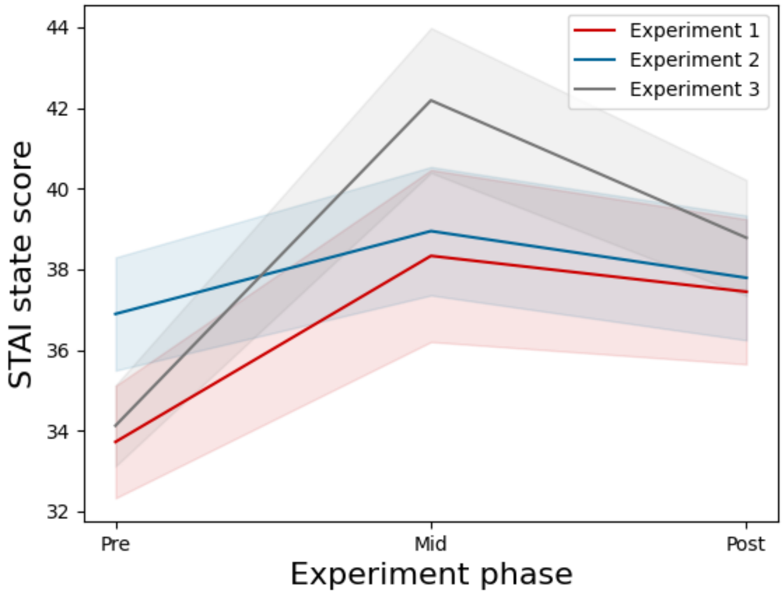


**Figure 1.** STAI state scores over time in Experiments 1, 2, and 3. Shaded areas represent the SEM.

The sleepiness of participants was measured after every other block, and showed significant changes over all experiments (Figure 2); Experiment 1, $F(5, 85) = 15.33$, $p < .001$, $\eta 2p = 0.47$, Experiment 2, $F(5, 90) = 7.93$, $p < .001$, $\eta 2p = 0.31$, and Experiment 3, $F(5, 155) = 32.78$, $p < .001$, $\eta 2p = 0.51$. Post-hoc comparisons revealed a significant increase between the pre-experiment sleepiness score, and all other scores in Experiment 1 and 2 (all $p < .001$), and no further significant differences. In Experiment 3, there was a similar significant increase between the pre-experiment score and subsequent scores (all $p < .001$), and a significant increase between sleepiness at block 2, and all blocks until block 8 (all $p < .01$), but no difference between block 2 and the sleepiness at the end of the experiment ($p = .08$). The decrease in sleepiness at block 8 and at the end of the experiment was also significant ($p = .008$).

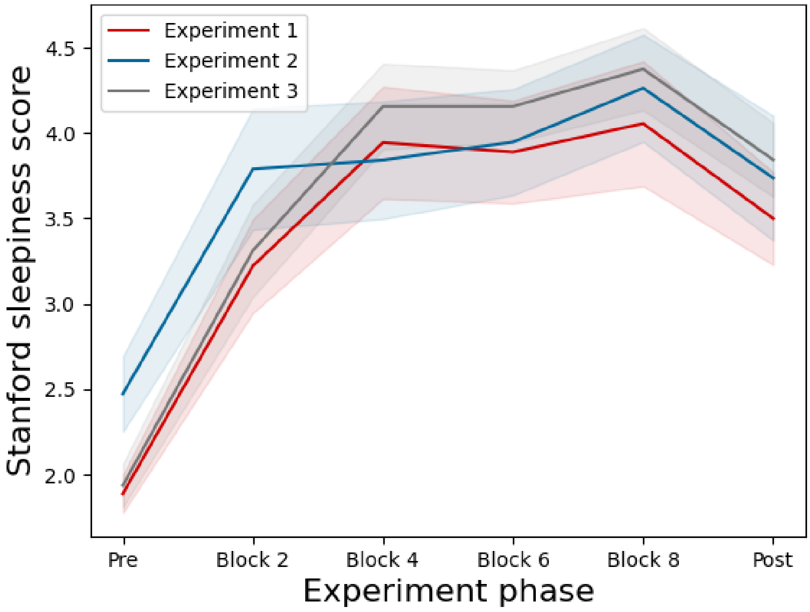


**Figure 2.** Stanford sleepiness scores over time in Experiments 1, 2, and 3. Shaded areas represent the SEM.

The positive and negative affect were measured before, during and after the experiment (Figure 3). The positive effect differed significantly in Experiment 1, $F(2, 34) = 12.43$, $p = .001$, $\eta 2p = 0.42$, which showed a decrease during the experiment compared to before ($p = .003$), but did not differ during or after the experiment. The negative affect in Experiment 1 showed no significant difference, $F(2, 34) = 1.19$, $p = .31$, $\eta 2p = 0.07$. In Experiment 2, there was a significant effect of positive affect as well, $F(2, 36) = 9.08$, $p < .001$, $\eta 2p = 0.34$. The positive affect decreased during the experiment ($p = .003$), but did not differ during or after the experiment ($p = .12$). The negative affect did not show a significant effect, $F(2, 36) = 0.60$, $p = .55$, $\eta 2p = 0.03$. In Experiment 3, there was an effect of positive affect, $F(2, 62) = 36.45$, $p < .001$, $\eta 2p = 0.54$, where the positive affect decreased during the experiment compared to before ($p < .001$), and also increased after the experiment compared to during the experiment ($p = .005$). The negative affect in Experiment 3 also differed significantly throughout the experiment, $F(2, 62) = 8.09$, $p = .003$, $\eta 2p = 0.21$. However, post-hoc comparisons revealed no significant changes when comparing the negative affect before and during the experiment (*p* = .012) or between during the experiment and after the experiment (*p* = .012).

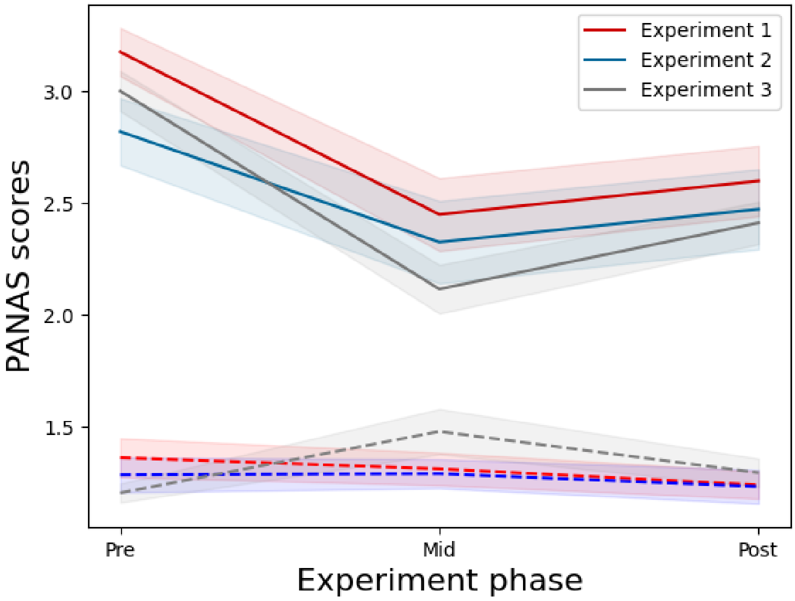


**Figure 3.** Positive affect (solid) and negative affect (dashed) scores over time in Experiments 1, 2, and 3. Shaded areas represent the SEM.
